# Trends in mosquito species distribution modeling: insights for vector surveillance and disease control

**DOI:** 10.1101/2023.03.16.533046

**Authors:** Catherine A. Lippi, Stephanie J. Mundis, Rachel Sippy, J. Matthew Flenniken, Anusha Chaudhary, Gavriella Hecht, Colin J. Carlson, Sadie J. Ryan

## Abstract

Species distribution modeling (SDM) has become an increasingly common approach to explore questions about ecology, geography, outbreak risk, and global change as they relate to infectious disease vectors. Here, we conducted a systematic review of the scientific literature, screening 563 abstracts and identifying 204 studies that used SDMs to produce distribution estimates for mosquito species. While the number of studies employing SDM methods has increased markedly over the past decade, the overwhelming majority used a single method (MaxEnt) and focused on human infectious disease vectors or their close relatives. The majority of regional models were developed for areas in Africa and Asia, while more localized modeling efforts were most common for North America and Europe. Findings from this study highlight gaps in taxonomic, geographic, and methodological foci of current SDM literature for mosquitoes that can guide future efforts to study the geography of mosquito-borne disease risk.

## Background

Mosquito-borne diseases have long imposed a heavy burden on both human and animal health worldwide [1]. There is an extensive history of mosquito control efforts to reduce the transmission of mosquito-borne diseases of global public health importance, notably malaria, yellow fever, and dengue fever [2–4]. However, these efforts are increasingly undermined by the combined effects of climate change, urbanization, and health system erosion, all of which are implicated in the expansion of mosquito-borne diseases to higher latitudes and elevations [3,5,6], the re-emergence of diseases like malaria and yellow fever [7–10], and the emergence of novel pathogens like chikungunya and Zika virus [11–14]. Despite their public health importance, most of these diseases are under-surveilled and under-reported, particularly in areas where poverty overlaps with a growing number of syndemic and syndromically hard-to-distinguish mosquito-borne diseases [15–17]. As a result, mapping the geographic distribution of mosquito vectors is often used as a first step towards describing the shifting landscapes of infectious disease risk.

One of the most commonly applied tools to study mosquito geographic distributions is species distribution modeling (SDM). Also commonly known as ecological niche modeling (ENM), species distribution models relate presence-absence or presence-only occurrence data to explanatory landscape factors, producing estimates of suitable habitat [18,19]. Inputs for SDMs typically include geolocated data on the presence of the species of interest as the response variable, often in the form of occurrence records derived from literature reviews, databases, or aggregated abundance sampling [18]. Explanatory variables are extracted from a wide range of sources, and ideally represent aspects of the species’ ecology that impact whether the organism can persist in a particular environment. Researchers often consider climatological factors, as well as place and organism-specific factors such as land cover, slope, aspect, elevation, soil type, and human effects on the landscape [20–22].

Given the flexibility of the approach, the motivations and objectives for developing SDMs of mosquito species often vary alongside model inputs, methods, and spatial scope. For example, global, regional, or national-scale SDMs may aim to anticipate broad distributions of present or future disease risk [23–25], while models developed at a finer spatial scale play an increasingly important role in vector control strategies. Public health vector control and mosquito source reduction are cornerstones of disease management, curbing transmission when clinical treatment and prophylactic options are limited or non-existent, as is currently the case for many arboviruses [26]. Identifying areas at risk from mosquito-borne disease transmission is integral to the development of effective policies, formation of mitigation strategies, and allocation of resources [27,28]; however, vector surveillance activities can be resource intensive and geographically limited [29,30]. Research-guided mosquito surveillance and abatement efforts are therefore often cited as necessary for practitioners to precisely know not only when, but also where to both monitor and intervene [31–37].

Here, we undertake a systematic review of the current body of literature on mosquito SDM research, motivated by the desire to understand these different applications, and to identify trends, challenges, and gaps in the current body of knowledge generated around mosquito ecology and biogeography. We followed the Preferred Reporting Items for Systematic Reviews and Meta-analyses (PRISMA) guidelines to identify and compile studies that developed SDMs of mosquito species in the past twenty years and extracted information on the species, methods, input variables, and location and scale of each relevant published, peer-reviewed study [38].

## Methods

We conducted literature searches following Preferred Reporting Items for Systematic Reviews and Meta-analyses (PRISMA) statement guidelines, a checklist of criteria to ensure transparency in systematic reviews [38,39]. We conducted searches across all Web of Science databases through January 2023 to identify studies using SDMs to estimate mosquito geographic distributions. Combinations of key terms used in searches included “Aedes”, “Culex”, “Anopheles”, and “mosquito” with “species distribution model*” and “ecological niche model*”. We included *Aedes, Anopheles*, and *Culex*, as explicit search terms, as these genera comprise the disease vectors most targeted in public health initiatives, and are intensely studied as disease vectors [40–42]. The inclusion of “mosquito” as a search term was to ensure that we captured studies on species beyond these three taxonomic groupings, and we did not restrict our searches to species implicated in human disease transmission. While there were no restrictions on the geographic region of study or date of publication, searches were limited to English language results.

Duplicate records were removed from our search results before screening. We screened the remaining abstracts for subject relevance (i.e., studies on mosquitoes), additionally removing publications that were literature reviews, expert commentaries, synthesis papers, phylogenetic studies, or gray literature. The remaining studies were reviewed in full for inclusion, excluding studies with methodologies that were not within the scope of review, including papers that were purely descriptive, used mosquito-borne disease cases as response variables, or modeled mosquito presence, abundance, or behaviors (e.g., such as oviposition or bite rates) using data with no geospatial component.

We extracted information from the full text of the remaining studies, which included information on publication (e.g., DOI link), the mosquito species of interest, SDM methods used, geographic location of study, spatial scale of analysis, and data sources for both species occurrence records and explanatory environmental variables. We noted methods for methods used for model fitting, addressing collinearity, and if modeled distributions were projected beyond their initial training scope (e.g., models projected to other geographic locations, or future time horizons) when available. Methods used in studies were classified into nine categories, outlined in Table 1.

**Table 1.** Nine categories of methods defined in this review that were used to build species distribution models.

| Modeling method | Definition (example) |
| --- | --- |
| CTA | **C**lassification **T**ree **A**nalysis methods, including classification and regression trees (CART), boosted regression trees (BRT), and random forest (RF) |
| Ensemble | A weighted or unweighted average, or combination, of models built with different methods |
| Envelope | Models that identify the boundaries of species' ecological tolerance directly from data, without the use of machine learning (e.g., BIOCLIM, CLIMEX, CliMond, DOMAIN) |
| GARP | **G**enetic **A**lgorithm for **R**ule-set **P**roduction, generates mathematical rules for estimating species presence |
| MaxEnt | **M**aximum **E**ntropy for expressing probability distributions |
| Mechanistic | Process-based models, often using parameters of physiological limits to estimate distributions (e.g., species thermal limits) |
| Mixed | Using two or more methods to estimate species distributions, but does not average or combine output into a model ensemble |
| Regression | Non-machine learning regression models (e.g., logistic regression, generalized linear models (GLM), generalized additive models (GAM), etc.) |
| Other | Less commonly used methods that did not fit in another category (e.g., ecological niche factor analysis (ENFA), environmental suitability thresholds, logic thresholds) |

Data extracted from the final collection of screened literature were synthesized to describe trends in mosquito SDMs. Data visualization was conducted in R (v4.1.2.) using code adapted from Lippi et al. 2021 [43], and mapping was performed in ArcMap (v10.8.1). The database of screened literature is available on GitHub (https://github.com/RyanLab/MOSQ_SDM_Table).

## Results

The initial search returned 1,185 records (Fig. 1), and 563 records remained after duplicates were removed. In initial abstract screening, 298 records for studies that did not fit the scope of this review were removed. After reviewing the full text of the remaining 265 records, we retained 204 studies that met our criteria for inclusion (https://github.com/RyanLab/MOSQ_SDM_Table).

**Figure 1.**
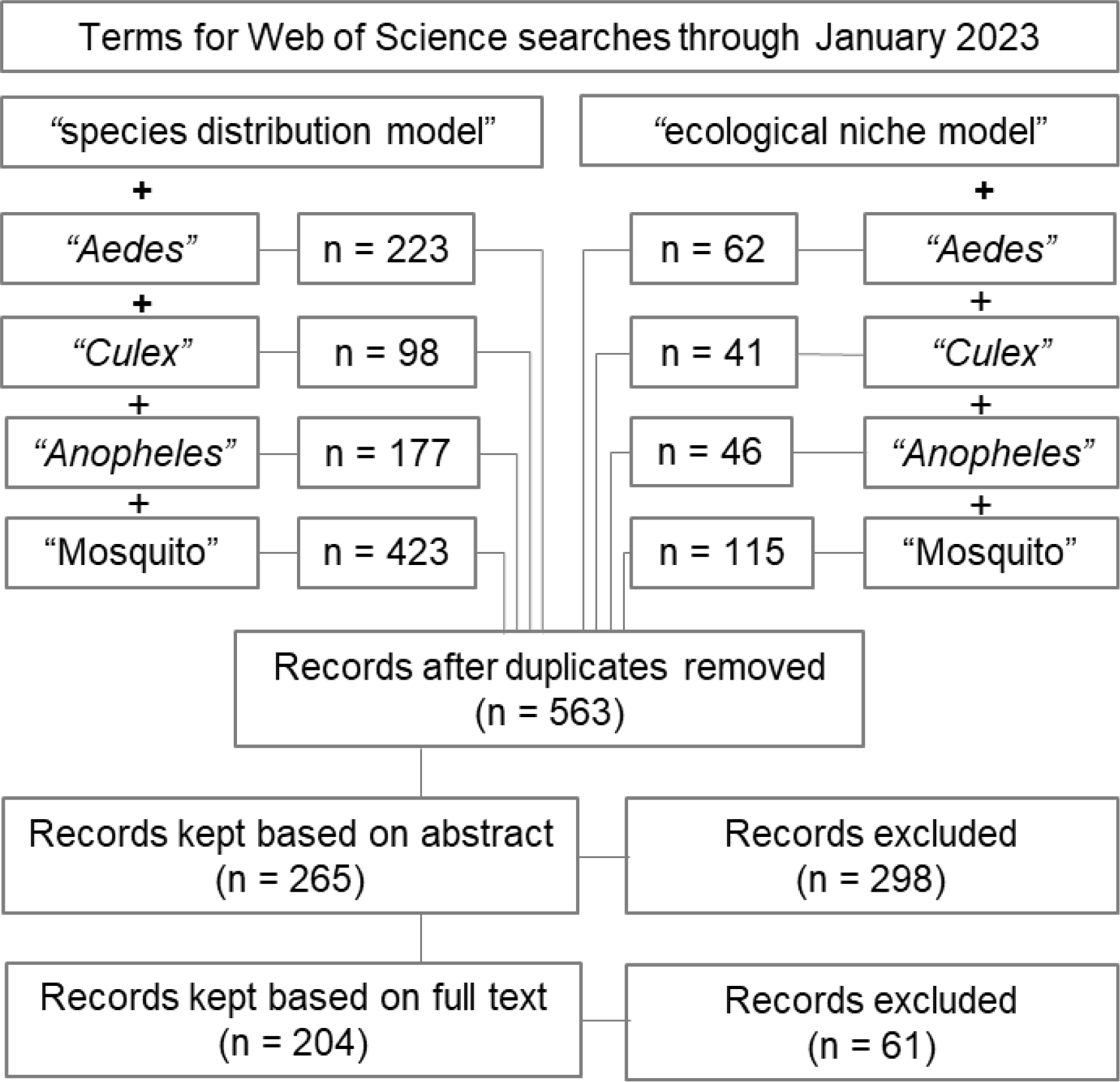
Flow diagram of the systematic review process, indicating combinations of search terms and number of studies screened.

### Taxonomic focus of SDMs

There were 138 mosquito species featured in SDMs produced in the reviewed literature, which included 78 species in *Anopheles*, 25 species in *Culex*, 24 species in *Aedes (= Ochlerotatus)*, and 11 species in other genera, including *Coquillettidia, Culiseta, Haemagogus*, and *Sabethes*. By species, most SDM studies developed models for *Ae. aegypti* (n = 55), *Ae. albopictus* (n = 50), *Cx. pipiens* (n = 20), *An. gambiae* (n = 17), and *An. arabiensis* (n = 15) (Fig. 2).

**Figure 2.**
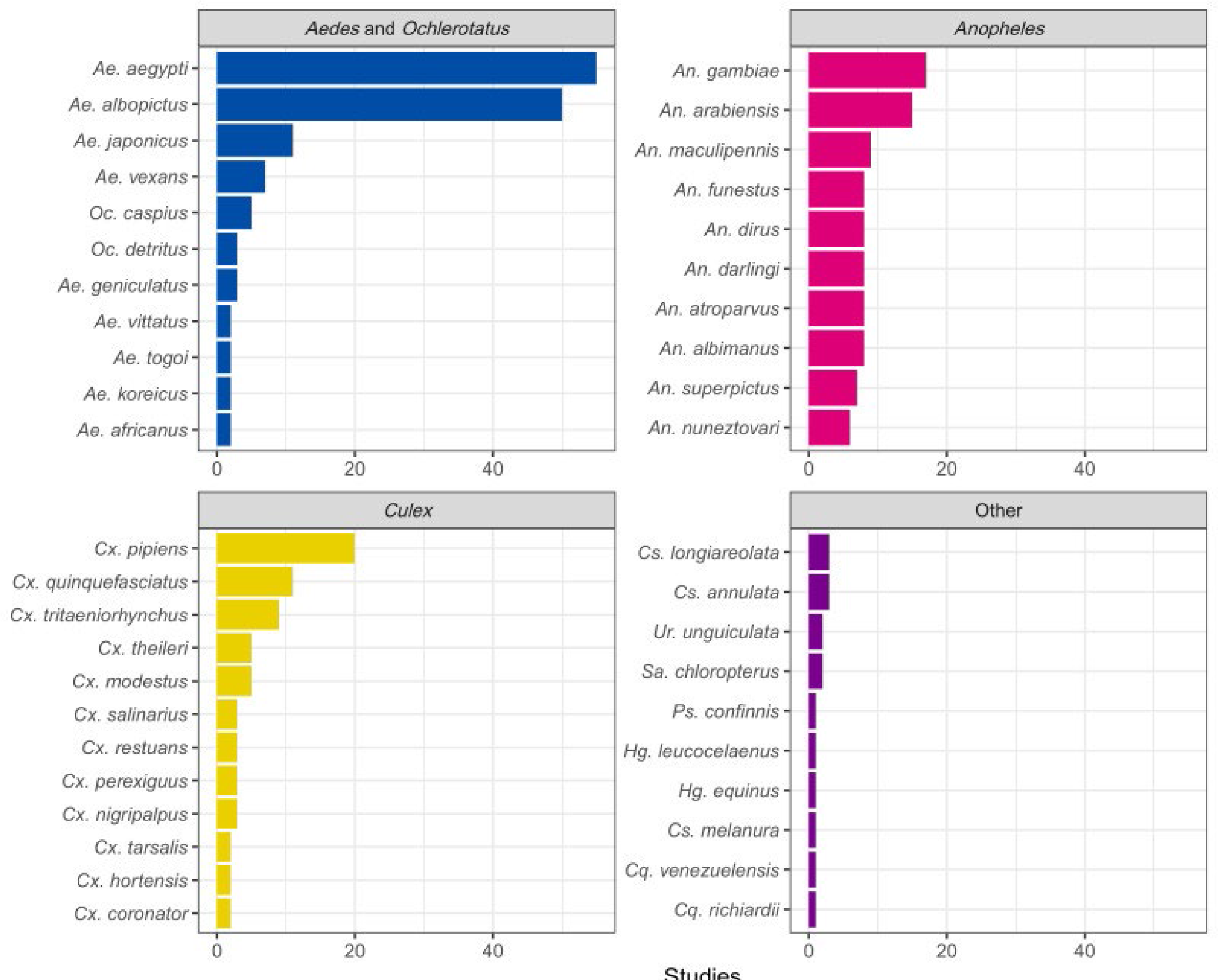
Top ten mosquito species, within each genus, that have been studied with species distribution models (SDMs). Most efforts focused on *Ae. aegypti* and *Ae. albopictus*, followed by *Cx. pipiens*, though collectively species in the genus *Anopheles* were also extensively modeled (*Cs. = Culiseta, Cq. = Coquillettidia, Hg. = Haemagogus, Ps. = Psorophora, Sa. = Sabethes, Ur*.*= Uranotaenia*).

Grouping by genera, mosquitoes featured in SDM studies have changed over time (Fig. 3A). The early mosquito SDM literature predominantly focused on mosquitoes in the genus *Anopheles*, comprising 64.3% of studies published through 2010. In recent years, studies on *Aedes* mosquitoes have become more prevalent, and were the taxonomic focus of nearly half (48.6%) of all mosquito SDMs produced since 2015. In the same period, roughly a quarter of studies were on mosquitoes in *Anopheles* (24.0%), and 11.6% in *Culex*.

**Figure 3.**
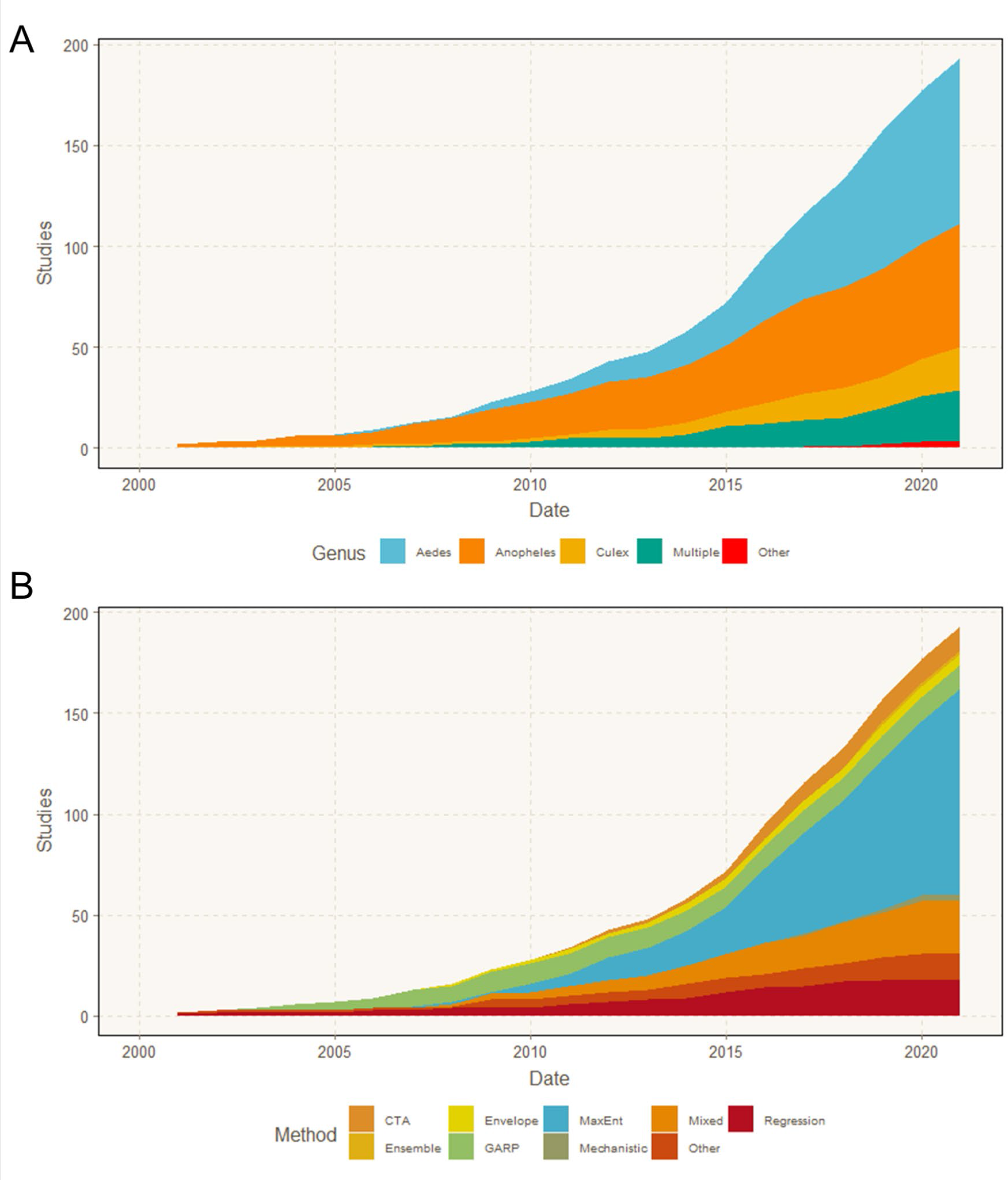
Cumulative number of studies on mosquitoes that modeled a given mosquito genus (A), and the SDM methods used in studies over time (B).

### Methods used to build SDMs

Overall, 169 of the 204 studies reviewed (82.8%) used a single method to estimate mosquito distributions, while the remaining 35 (17.2%) used more than one method or used model ensemble approaches (Fig. 3B). More than half (n = 108, 52.9%) of the studies used MaxEnt exclusively, a trend which also holds across genera (Fig. 4). An additional 21 (10.3%) used MaxEnt with one or more additional SDM methods. Non-machine learning regression models (e.g., logistic regression, generalized linear models, etc.) were used exclusively in 18 studies (8.8%) and used in combination with other methods in an additional 16 studies (7.8%). Classification tree analysis (CTA) methods, which included classification and regression trees (CART), boosted regression trees (BRT), and random forest (RF), were used as the sole SDM method in 14 studies (6.9%), and the Genetic Algorithm for Rule Set Prediction (GARP) was the sole method in 12 studies (5.9%). Classification tree analysis methods were combined with other SDMs in 18 additional studies (8.8%), while GARP was used with other methods in 5 additional studies (2.5%). Mechanistic models were used as the only SDM method in 3 studies (1.5%), and bioclimatic envelope models were used in 5 studies (2.5%). Thirteen studies (6.4%) were characterized as “Other,” featuring uncommon methods such as ecological niche factor analysis (ENFA), and other types of environmental suitability or logic thresholds.

**Figure 4.**
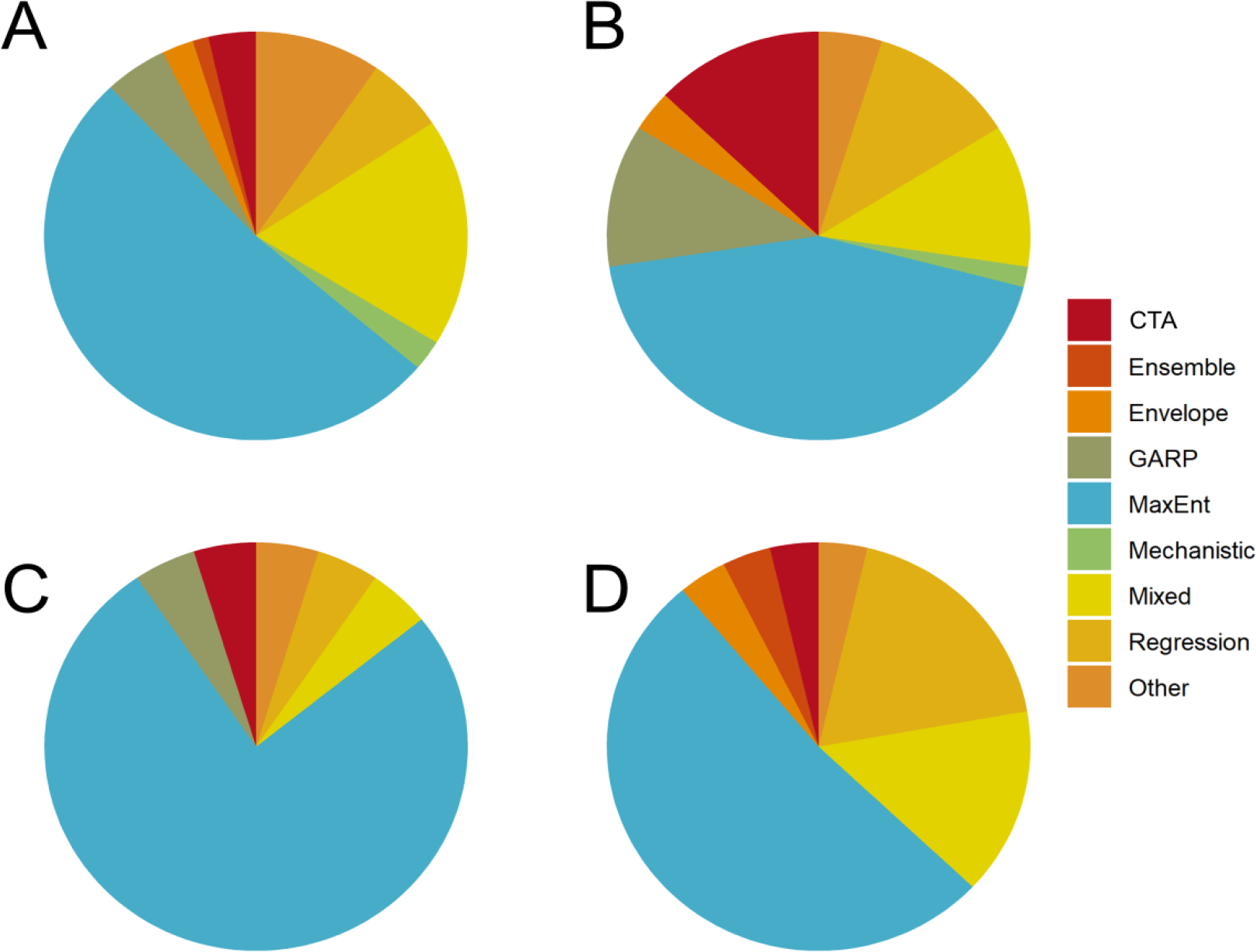
Breakdowns of methods used, shown by taxonomic groups for *Aedes* (A), *Anopheles* (B), *Culex* (C), and studies that estimated distributions for multiple genera (D).

The frequency of using SDMs to estimate mosquito distributions increased markedly over time, particularly from 2015 onward (Fig. 3B). Concurrently, the evolution of SDM algorithms led to a more diverse methodological landscape. In the early 2000’s, GARP was the most commonly implemented method among these studies, and remained a frequently used method until approximately 2010. MaxEnt software was released in 2006, and the first mosquito SDM study using MaxEnt was published shortly thereafter in 2007. By 2011, it was the most common SDM approach, with 19 of 43 papers (44.2%) during those years solely using MaxEnt. More than half (62.3%) of the mosquito SDM studies published since 2015 solely used MaxEnt to estimate mosquito distributions. To a lesser extent, the frequency of using multiple SDM approaches in a single study (“Mixed” in Figures 3 and 4) has also increased over time. The first mixed methods study was published in 2008, and studies that used multiple SDM approaches have accounted for 13.7% of those published since 2015. Classification tree analyses (CTAs) and regression methods have seen modest increases in use over the past decade, and were used in 8.2% and 6.2% of studies, respectively, since 2015. The increased availability of gridded data layers of ecological and climate products, representing a host of environmental factors, has also been a fundamental piece in the rapid expansion of SDM research. A notable example is the WorldClim database, which was first released in 2005 (with version 2 released in 2017) and made long-term averages of historical and projected future climate data accessible for many SDM studies [44].

### Spatial scale of SDMs

Scale of analysis varied considerably throughout the mosquito SDM literature. The majority of studies were conducted at the sub-national (34.8%) or national (31.4%) level. Most national or sub-national studies were conducted in the United States (11.1%), followed by Germany (6.7%), Mexico (6.7%), Australia (5.2%), Brazil (4.4%), China (4.4%), Colombia (4.4%), Italy (4.4%), Argentina (3.7%), Iran (3.7%), and Tanzania (3.7%) (Fig. 5). Approximately one quarter of studies (24.5%) were regional, projecting models over large geographic areas that encompassed multiple countries. Most of the regional models were developed for portions of Africa (30.0%) and Europe (30.0%), followed by SDMs developed for regions in Asia (24.0%), North America (16.0%), and South America (16.0%). Nearly half of all regional models (44.0%) were developed for species of *Anopheles* mosquitoes. Relatively few SDMs (9.3%) focused on a global extent, but of those, the majority (68.4%) focused on mosquitoes in the genus *Aedes* (Fig. 6A). Studies in African countries typically modeled the distributions of *Anopheles*, while European studies tend to focus on *Aedes*. In comparison, studies conducted in North America, South America, the Middle East, and countries in Asia more frequently included multiple species, or other taxonomic groups (Fig. 5).

**Figure 5.**
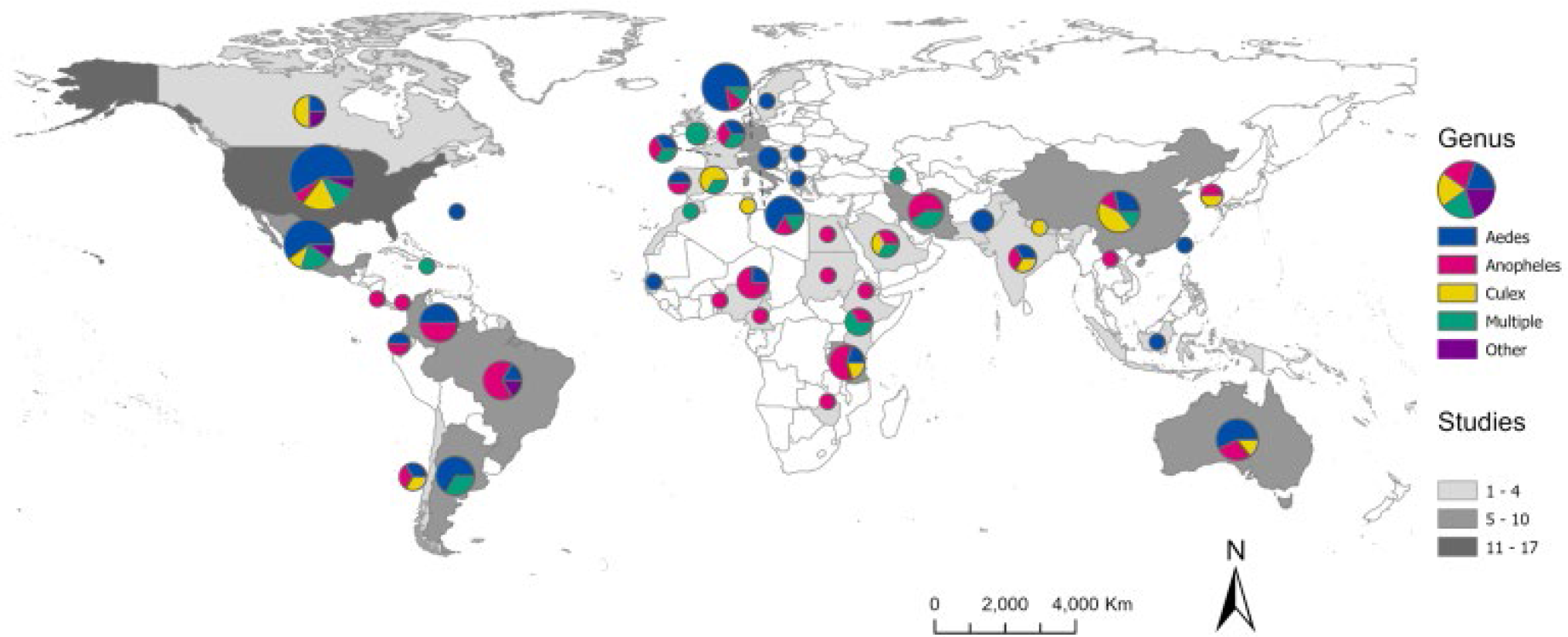
Map representing the number of SDM studies conducted at the national or sub-national level (country greyscale) and the mosquito genera modeled in the studies (pie chart color breakdown).

**Figure 6.**
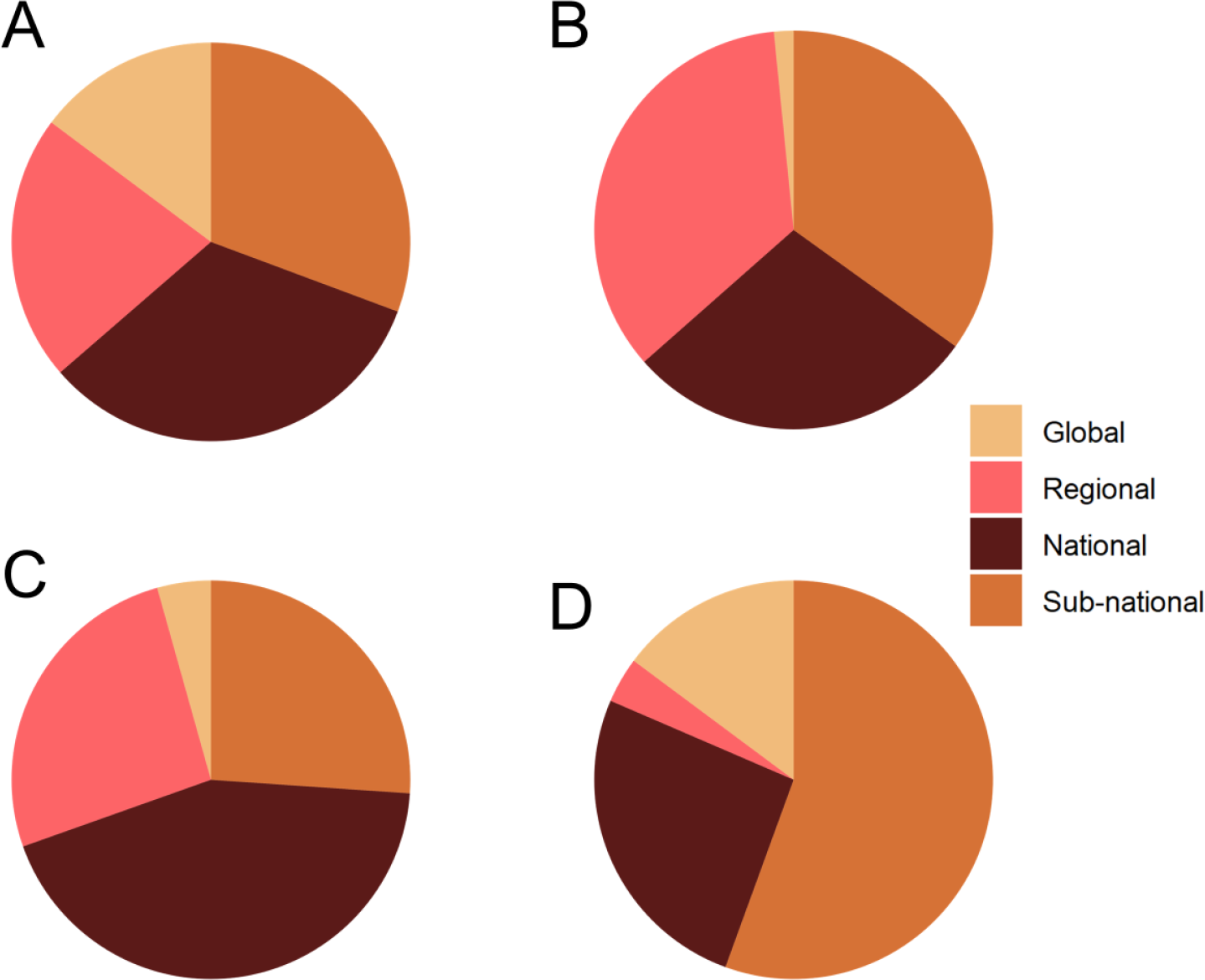
The proportion of spatial scales represented in the reviewed SDM studies, shown by genera for *Aedes* (A), *Anopheles* (B), *Culex* (C), and multiple genera (D).

The scale of analysis used to build SDMs also varied by taxa. Species distribution models were built for *Aedes* mosquitoes in roughly equivalent proportions (Fig. 6A), possibly driven by the global invasions of *Ae. aegypti* and *Ae. albopictus*. In contrast, global suitability models for *Anopheles* and *Culex* were relatively uncommon (Fig. 6B & 6C), regional and sub-national models were most common for *Anopheles* (Fig. 6B), and SDMs for *Culex* and multiple genera primarily consisted of national and sub-national models (Fig. 6C & 6D).

### Data sources used to build SDMs

Mosquito occurrence records used as data inputs for SDM workflows were obtained from a variety of sources. Over one third of the reviewed studies (36.8%) included georeferenced locations from online data repositories when building models, the majority of which (49.3%) used the Global Bioinformatics Information Facility (GBIF) to obtain data. Georeferenced data obtained from published sources and literature reviews were also frequently used as data inputs in studies (36.8%), with the database published by Kraemer et al. [45] explicitly cited as a data source in 5.4% of all studies. Use of existing public health surveillance systems for mosquito records, which included databases from national public health authorities, was indicated in 11.8% of studies.

Collection of novel mosquito records through entomological sampling was indicated in approximately one third (32.2%) of published studies, of which 16.1% supplemented collection records with data from other sources (e.g., published data, online repositories, etc.). Of the studies that collected entomological data, 59.3% reported larval sampling, of which 18.8% reported sampling with oviposition or gravid traps. Diverse sampling methods were reported in studies with entomological data that targeted the adult life stage, including CDC light traps (37.8%), BG-Sentinel traps (22.2%), Mosquito Magnet traps (15.6%), aspiration (13.3%), and human landing catch (11.1%). While studies that included a field component typically described the mosquito life stage targeted in sampling, across all studies the life stage of mosquitoes used for species presence (i.e., adults, immatures, or both) was not specified in most instances (53.9%).

### Environmental predictors of mosquito presence

Most studies (89.7%) incorporated climate variables to estimate mosquito distributions. While some studies (20.1%) used climate data exclusively, many (69.6%) used a combination of climate data and other environmental covariates (e.g., land cover class, elevation, soil classifications, etc.) in their model predictions. Most studies (69.2%) that used climate data to produce SDMs employed WorldClim data products. Many studies (27.9%) projected models to estimate mosquito distributions under future climate conditions, using products such as downscaled global climate models (GCMs) as environmental predictors. A variety of methods were used to control for collinearity in environmental predictors before building SDMs, including use of correlation coefficients with a threshold (27.5%), principal component analysis (PCA) (8.3%), and variance inflation factor (VIF) (5.9%). Nearly half of studies (48.5%) did not explicitly address predictor collinearity.

Variables identified as important for predicting mosquito distributions were reported in 80.3% of studies. Identified drivers varied considerably between studies and taxa. Measures of temperature were most often described in study text as important predictors of mosquitoes (54.9%), followed by precipitation (42.6%), land cover and land use (31.4%), and elevation (18.6%). Of the studies that incorporated climate variables, 62.6% identified temperature, and 52.5% identified precipitation as important predictors of mosquito habitat suitability. By taxonomic group, temperature variables were top predictors in 44 (50.0%) studies on *Aedes*, 37 (58.7%) studies on *Anopheles*, and 16 (69.6%) studies on *Culex*. Precipitation variables were top predictors in 32 (36.4%) studies on *Aedes*, 36 (57.1%) studies on *Anopheles*, and 12 (52.2%) studies on *Culex*. Variable importance was not reported in 19.6% of studies.

## Conclusions

Species distribution modeling has become a frequently used methodological approach to estimate the distribution, and implicit risk, of vector-borne diseases [35,36]. In this study, we conducted a systematic review of scientific literature that used SDMs to estimate geographic distributions of mosquitoes. By quantifying data from the screened literature to identify patterns and trends, we were able to summarize the methods, taxonomic foci, geographic scope, and other attributes reported in SDM studies. Importantly, this also enabled us to identify potential gaps in the current literature, providing guidance for future modeling efforts.

### Current trends in mosquito SDMs

Although there is diversity in the landscape of available modeling approaches and tools, MaxEnt is the most commonly implemented method for conducting SDM studies on mosquitoes [46]. After its release in 2006, MaxEnt quickly gained favor over previously common methods, like GARP, and remains the most frequently used approach used to estimate mosquito distributions [47]. While MaxEnt may be the most appropriate methodology for some studies, the popularity of the method also results from an interpretable graphic user interface, prolific training guides, and general ease of implementation for users. Further, the release of WorldClim climate model output data in 2005, and the subsequent availability of other gridded environmental data products, allowed users to perform analyses without collecting primary environmental data, facilitating the use of SDMs [48].

The majority of the SDM studies in this review were for *Anopheles* or *Aedes*. This is not surprising, given the emphasis of global health importance of malaria transmitted by mosquitoes in the genus *Anopheles*, and arboviral pathogens transmitted by some mosquitoes in the genus *Aedes*, including YFV and DENV. Moreover, most global modeling efforts involve mosquitoes in the genus *Aedes*, likely owing not only to the medical importance of two key species (*Ae. aegypti* and *Ae. albopictus*), but also to their cosmopolitan success as invasive species. In contrast, SDMs for *Anopheles* were typically conducted at regional, national, and local scales, and in locations predominantly on the African continent, reflecting disproportionate research effort focused on *Plasmodium falciparum* malaria.

Geographic gaps exist for highly localized studies in Central Asia, Southeast Asia, Eastern Europe, and portions of Africa and South America. Notably, many of these regions have rich mosquito diversity, including dozens of understudied vectors of current or potential future emerging infections [49]. Modeling studies in these regions were also underdeveloped in other methodological aspects; for example, temperature and precipitation in these regions were most commonly represented by WorldClim bioclimatic variables, as opposed to bespoke regional climate products. Regionally targeted research efforts may benefit from locally-created and locally-validated climate and land cover products.

### Challenges of SDMs

We found considerable variation in which environmental drivers were identified as predictive of mosquito distributions, such that few generalizations could be made even for a given species. Variable importance is influenced by nearly every step of the SDM building process, such as choice of data products, scale of analysis, collinearity reduction techniques, and choice of SDM algorithm. In studies that reported variable importance, actual values of environmental predictors (i.e., numerical thresholds for occurrence) were rarely reported. The prevalence of studies that failed to reduce variables to account for collinearity, or lacked justification for choice of environmental predictors, points to a potentially troubling neglect of biological grounding and hypothesis testing. These challenges can be readily addressed in future studies through adherence to best practices and standards in building models and reporting results [50–52]. Assessing the quality of models and adherence to best practices is beyond the scope of this review, but has been recently assessed by Barker and MacIsaac [51]. Given some of these underlying heterogeneities among studies, care must be taken when interpreting the results of SDMs, especially those potentially used for guiding public health decisions, as poor quality models can divert resources and miscommunicate the true risk of exposure.

### Opportunities for future modeling efforts

Many efforts have been made to delineate the geographic extent of mosquitoes, but the current literature still may not capture the full landscape of risk, especially in the biodiverse areas where new infections are more likely to originate [53,54]. The movement of vectors and pathogens may also serve as the catalyst for new public health challenges, for example, when mosquito vectors aggressively invade new locations [6,55], or when the introduction of pathogens increases the medical significance of local mosquito populations [35]. These changes also create a problem for the scientific literature itself: estimation of range boundaries based on baseline climate conditions has diminishing value in a rapidly-warming world, where mosquito ranges have already become non-stationary in both invasive ranges [25] and endemic areas [6].

Our findings therefore indicate an opportunity not only to expand data collection and distribution modeling efforts for under-represented mosquito species and in underrepresented areas [43,56], but to more broadly rethink the SDM workflow as it is currently used in vector surveillance and control. An iterative workflow is technologically feasible and cost-effective, where (1) existing surveillance data and local knowledge are used to generate or update mosquito distribution models; (2) new forecasts are generated that anticipate areas at risk of range expansions, based on existing trends and climate projections; and (3) models are used to guide the collection of new surveillance data (Fig. 7). Field efforts to collect data used for model improvement can be a daunting endeavor, yet we found a surprisingly high number of studies which collected novel entomological survey data for building models. Nevertheless, relatively few studies incorporated data from public health surveillance systems, highlighting a potential avenue to future collaborations between modelers, public health authorities, and vector control agencies. This new workflow presents opportunities on the technological front, where efforts to employ newer approaches with updated methodologies and software, and adherence to best practices, may enable us to refine estimates of spatial risk; moreover, adoption of automated approaches that update range estimates from incoming data could aid in making timely predictions that are more accessible to decision makers. Partnerships with local experts and agencies will be key both to improving model predictions and maximizing their applied utility [57]. Ultimately, working more closely with end users may facilitate the uptake of modeling workflows, ensuring that SDMs are appropriately contextualized and regularly updated.

**Figure 7.**
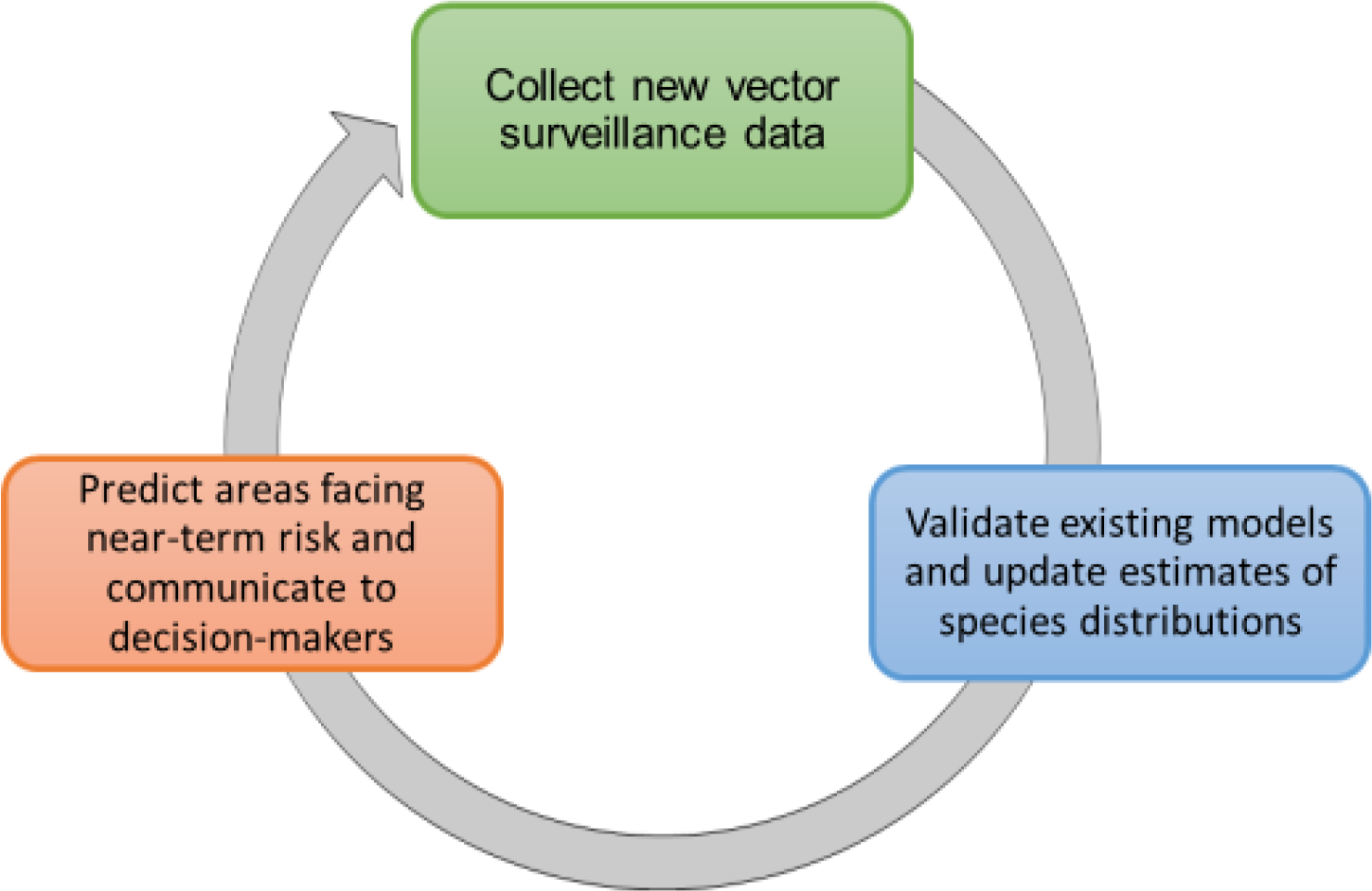
A conceptual workflow for dynamic mosquito species distribution modeling.

## List of Abbreviations

BRT: boosted regression trees
CART: classification and regression trees
CTA: classification tree analysis
DENV: dengue virus
ENM: ecological niche model
GBIF: Global Bioinformatics Information Facility
GCM: global climate model
GLM: generalized linear model
PCA: principal component analysis
RF: random forest
SDM: species distribution model
VIF: variance inflation factor
YFV: yellow fever virus

## Declarations Ethics approval and consent to participate

Not applicable

## Consent for publication

Not applicable

## Availability of data and material

Data associated with this analysis are available on GitHub (https://github.com/RyanLab/MOSQ_SDM_Table).

## Authors’ contributions

Conceptualization: CAL, SJM, and SJR; Analysis: CAL, SJM, RS, MF, AC, GH; Visualizations: CAL, CJC, SJR; First draft writing: CAL, SJM, SJR, RS; Final draft and reviewing: all authors

## Acknowledgements

We would like to thank members of the QDEC lab for conversations and company while creating and revising this paper.

